# Pharmacological reduction of neutrophil infiltration reduces *Clostridioides difficile* infection severity

**DOI:** 10.1101/2025.10.30.685526

**Authors:** Orlaith Keenan, Joshua Soto Ocaña, Alexa Semon, Tiffany H. Zhou, Emma E. Furth, Gavyn Chern Wei Bee, Daniel L. Aldridge, Juliana Diamantino, Christopher A. Hunter, Ken Cadwell, David M. Aronoff, Joseph P. Zackular

## Abstract

*Clostridioides difficile* is the leading cause of nosocomial infections and an urgent public health threat. This bacterial pathogen is challenging to treat due to antibiotic resistance and high recurrence rates, highlighting the need for additional therapeutic strategies. The host inflammatory response is a major driver of *C. difficile*-associated disease and associated with worse clinical outcomes. Currently, few strategies targeting the inflammatory response have been leveraged to treat CDI. Here, we show that administration of the prostaglandin E_1_ (PGE_1_) analog misoprostol markedly reduces CDI severity by modulating host immune responses. During CDI, misoprostol decreases circulating neutrophils and limits infiltration into the colon, reducing epithelial damage, intestinal pathology, and infection severity. Additionally, misoprostol reduces serum granulocyte colony-stimulating factor (G-CSF), an important cytokine in neutrophil mobilization, controlling neutrophil levels during CDI. Together, these findings highlight neutrophil infiltration as a key driver of *C. difficile-*associated disease and identify innate immune modulation as a potential host-directed therapeutic strategy.

## INTRODUCTION

*Clostridioides difficile* is a leading cause of antibiotic-associated diarrhea and remains a major public health threat worldwide(1, 2). *C. difficile* causes a wide range of disease outcomes and is challenging to treat due to high rates of antibiotic resistance, recurrence, and limited therapeutic options. Antibiotic use is the primary risk factor for *C. difficile* infection (CDI), as it disrupts the resident microbiota, allowing *C. difficile* spores to germinate and produce toxins that damage the intestinal epithelium and trigger a strong inflammatory response(3). This inflammation correlates strongly with CDI severity(4, 5), highlighting the central role of the immune response in shaping CDI pathology. Although antibiotics remain the leading treatment, they fail to resolve toxin-induced inflammation, and current immunomodulatory therapeutic efforts, including monoclonal antibodies and vaccines, primarily focus on neutralizing toxins(6, 7). While these strategies reduce CDI severity, they do not address the underlying inflammatory processes that drive pathology during acute CDI, highlighting the need for host-directed approaches that modulate immune responses to improve CDI outcomes.

The innate immune response plays a pivotal role in shaping infection outcomes(4, 5). Following toxin-mediated damage to the intestinal epithelium, neutrophils and monocytes are rapidly recruited to the gut, where they release cytokines and chemokines that amplify inflammation and recruit additional immune cells(8). Neutrophil influx into the epithelium and submucosa is a key marker of CDI and can be both protective and pathogenic(9-11). While neutrophils are important for controlling infection, they also release inflammatory molecules, including reactive oxygen species (ROS), proteases, and neutrophil extracellular traps (NETs), that can damage the epithelium and impair wound healing in colitis(12, 13). Neutrophil depletion in murine models of CDI demonstrates that these cells are necessary for host survival during acute infection(9, 11). However, excessive neutrophil recruitment can exacerbate pathology during CDI(10) and also contribute to the formation of pseudomembranes, a feature associated with severe *C. difficile* colitis(14). These findings highlight the importance of a balanced neutrophil response during infection; however, how this may be achieved is still unclear. A more complete understanding of neutrophil biology during CDI could thus yield promising targets for therapeutic intervention.

Prostaglandins (PGs) are short-lived lipid mediators produced through the arachidonic acid pathway that play important roles in maintaining intestinal homeostasis and regulating inflammation(15). We previously reported that misoprostol, an FDA-approved stable PGE_1_ analog that pharmacologically mimics the endogenous molecule PGE_2_(16), protects mice from CDI by improving survival, reducing intestinal permeability, and promoting recovery of the microbiota following antibiotic perturbation(17). However, the mechanisms underlying this protection remain unclear. While misoprostol has known cytoprotective effects in the gastrointestinal tract(18), whether it acts by modulating *C. difficile* virulence, enhancing epithelial barrier integrity, or regulating host immune responses during CDI has not been defined. PGE_2_, the most abundant PG, has well described roles in limiting intestinal inflammation and exerts context-specific effects on epithelial homeostasis and immune function in the gut(19-22). During colitis, PGE_2_ promotes epithelial proliferation, enhances mucus production, and modulates proinflammatory cytokine expression to support intestinal barrier integrity(23-25). Similarly, during intestinal *Toxoplasma gondii* infection, PGE_2_ limits proinflammatory cytokine production and neutrophil activation in the small intestine(19). While the roles of PGs during CDI remain poorly understood, these findings suggest that PGs may restrict inflammation and promote mucosal repair during CDI.

In this study, we demonstrate that misoprostol mitigates CDI severity by improving intestinal barrier function and modulating host inflammatory responses. We report that misoprostol protects against severe CDI by reducing circulating neutrophils in the blood and limiting their influx into the colon. We further show that misoprostol controls neutrophil levels during CDI by reducing levels of granulocyte colony-stimulating factor (G-CSF), an important cytokine in neutrophil development and mobilization(26), in peripheral blood. Conversely, exogenous supplementation of G-CSF attenuates the protective effects of this treatment. These findings provide new insight into the contribution of neutrophils to *C. difficile* pathogenesis and highlight how pharmacologic modulation of the host response can be leveraged to develop effective host-directed therapies for CDI.

## RESULTS

### Misoprostol reduces CDI severity by limiting colonic inflammation and epithelial injury

Pharmacologic modulation of host immune responses during CDI remains a promising avenue for treatment of severe acute infection. Our previous work demonstrated that misoprostol has therapeutic potential for CDI, rescuing mice from lethal infection(17). However, the mechanisms underlying this protection remain unclear. To further define mechanisms of misoprostol’s protective effects during acute CDI, we leveraged a mouse model of infection in which misoprostol was administered daily by intraperitoneal (i.p.) injection beginning one day prior to infection with *C. difficile* (Fig. 1A). Consistent with previous findings, misoprostol-treated mice (Cd + miso) exhibited less severe disease compared to untreated *C. difficile*-infected mice (Cd), as measured by a composite score that evaluates weight loss, stool consistency, and behavior(27) (Fig. 1B). Next, we sought to define the mechanism of misoprostol-mediated protection during infection. The impact of misoprostol on pathogen dynamics has not been established, so we first measured *C. difficile* burdens and toxin production. Notably, misoprostol treatment did not affect the abundance of *C. difficile* or stool titers of toxin throughout the infection (Fig. 1C-D), suggesting that PG treatment does not impact *C. difficile* virulence and instead, likely modulates the host response to reduce CDI severity.

**Fig. 1.**
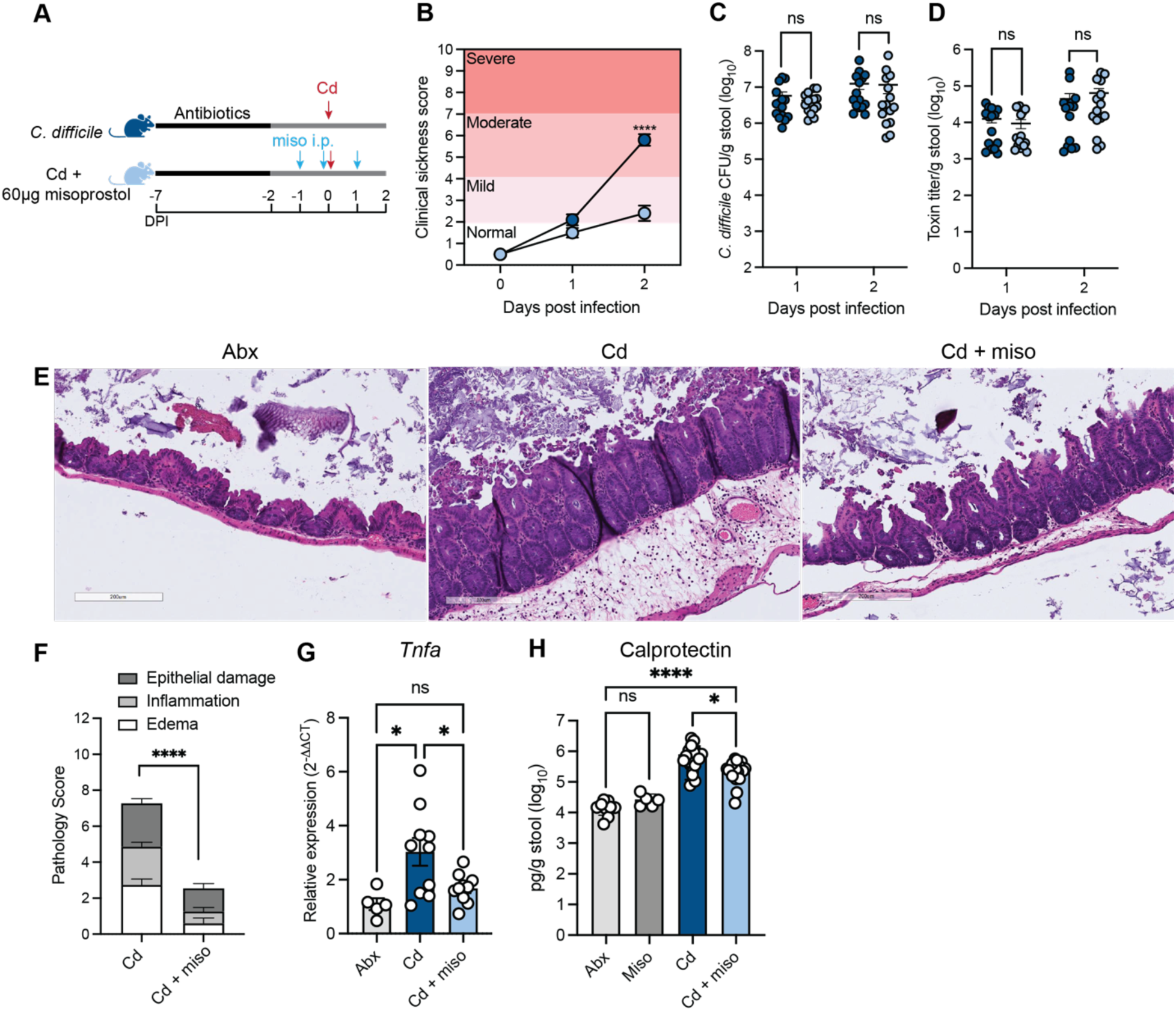
Misoprostol reduces CDI severity by limiting colonic inflammation and epithelial damage. (**A**) Schematic of experimental design. Mice were treated with cefoperazone (Abx) in their drinking water and infected with 1x10^5^ *C. difficile* spores (Cd). Misoprostol-treated mice (Miso or Cd + miso) were administered 60μg misoprostol via intraperitoneal (i.p.) injection daily from -1 days post-infection (DPI) to 1 DPI. (**B**) Composite score of weight loss, stool consistency, and behavior (N = 15 per group). (**C**) *C. difficile* colony-forming units (CFU) and (**D**) toxin titers from the stool of infected mice at 2 DPI (N = 15 per group). (**E**) Representative images of H&E-stained cecal slides from infected mice at 2 DPI. (**F**) Pathology scores from H&E-stained cecal slides at 2 DPI (N = 15 per group). (**G**) RT-qPCR on bulk colon tissue harvested from mice at 2 DPI. Data relative to average *Gapdh* expression of uninfected antibiotic-treated mice (Abx) (N = 5 – 10 per group). (**H**) Fecal calprotectin levels measured by ELISA at 2 DPI. (N = 5 – 15 per group). (**B**–**H**) Data represented as mean ± SEM and representative of at least two independent experiments. Statistics by two-way ANOVA with Sidak’s multiple comparisons (B), Mann-Whitney multiple comparison’s test (C - D), Welch’s unpaired t test (F), and one-way ANOVA with Tukey’s multiple comparisons (G) or Dunnett’s T3 multiple comparisons (H). ns (not significant) P>0.05, *P<0.05, ****P<0.0001.

Based on these findings and given the role of PGs as known immunomodulators(15, 21, 22), we postulated that the protective effects of misoprostol were through modulation of the host response to infection. To assess this, we first quantified intestinal pathology of H&E-stained cecal sections and found that misoprostol treatment significantly reduced histological markers of CDI severity, including epithelial damage, edema, and immune cell infiltration during CDI (Fig. 1E-F). Consistent with reduced pathology, Cd + miso mice also exhibited reduced canonical markers of inflammation, including reduced colonic expression of the inflammatory cytokine *Tnfa*, restoring levels to those observed in uninfected antibiotic-treated controls (Abx) (Fig. 1G). Additionally, levels of fecal calprotectin, a neutrophil-derived protein and marker of intestinal inflammation(28), were significantly reduced in Cd + miso mice compared to Cd mice (Fig. 1H).

The intestinal epithelium serves as one of the first lines of defense against CDI as *C. difficile* toxins damage epithelial cells, leading to a loss of barrier integrity, release of proinflammatory cytokines, and immune cell infiltration(8). Numerous studies have shown that PGs fortify the intestinal epithelium and promote recovery after damage(20, 21, 23). Given our previous finding that misoprostol treatment reduces intestinal barrier permeability during CDI(17) and our data showing reduced epithelial damage (Fig. 1F), we examined whether protection by misoprostol is mediated through effects on the epithelium. Lipocalin-2 is a metal sequestering protein that is produced by epithelial cells and serves as a marker of epithelial inflammation(29). Given its role in epithelial responses, we measured the levels of lipocalin-2 at two days post-infection. We observed no significant differences in lipocalin-2 levels during CDI following misoprostol treatment (Fig. S1A). We next measured canonical markers of inflammation in isolated intestinal epithelial cells (IECs) at two days post-infection(30). Interestingly, we found no significant differences in expression of *Tnfa* or the monocyte- and neutrophil-recruiting chemokines *Ccl2* and *Cxcl1*(31, 32). However, we did observe a trend toward decreased IEC expression of *Cxcl5*, a key neutrophil chemokine at mucosal sites(33) (Fig. S1B).

Based on the multifaceted roles that PGs play in shaping IEC responses, we examined whether misoprostol treatment during CDI impacts any of these key pathways(21, 23-25). One potential mechanism of protection is that misoprostol promotes expression of *Muc2*, a key mucin produced by epithelial cells that fortifies the physical barrier between IECs and the intestinal microbiota and is decreased in CDI patients(34). To test whether misoprostol impacts *Muc2* during CDI, we quantified expression in IECs at two days post-infection. Misoprostol treatment did not alter *Muc2* expression (Fig. S1C), suggesting that protection is not mediated through enhanced mucus production. Another potential mechanism of protection is that misoprostol promotes IEC proliferation during CDI which can fortify the intestinal barrier. To test this, we quantified Ki67-positive IECs within the intestinal crypts in stained cecal sections at two days post-infection, as Ki67 is an established proliferation marker(35). Consistent with prior PG studies(23), misoprostol-treated uninfected mice (Miso) showed increased Ki67 staining compared to Abx mice. However, we did not observe any differences in epithelial proliferation between Cd and Cd + miso mice (Fig. S1D-E). Collectively, these data support a therapeutic role for misoprostol and indicate that its protective effects during CDI are not mediated through changes in *C. difficile* virulence, mucus production, or IEC proliferation. Instead, these findings suggest that misoprostol confers protection by modulating the host immune response to infection.

### Misoprostol reduces colonic monocyte and neutrophil infiltration during CDI

Our data demonstrate that misoprostol is anti-inflammatory, decreasing immune cell infiltration and levels of the neutrophil-derived protein calprotectin during CDI (Fig. 1F, H). Neutrophils can exacerbate tissue damage during CDI(10); however, they are also critical in controlling disease as depletion of these cells increases CDI mortality(9). While the impacts of PGs on immune cell infiltration during CDI are unknown, PGE_2_ has been shown to reduce neutrophil infiltration into the lung during LPS-induced inflammation(36). Given these data and the important role of innate immune cell recruitment during infection, we postulated that misoprostol may modulate immune cell infiltration during CDI. To test this, we immunoprofiled the innate immune response during CDI in the colonic lamina propria (cLP) at two days post-infection(19) (Fig. S3A). We found that misoprostol treatment significantly reduced the total numbers of monocytes and neutrophils in the cLP (Fig. 2A-C). Next, to determine if misoprostol treatment affected expression of chemokines important for monocyte and neutrophil recruitment, including CCL2, CXCL1, CXCL2, CXCL5 (31, 32), we quantified chemokine gene expression in colonic tissues. We observed significant reductions in the expression of *Ccl2* and *Cxcl5* in Cd + miso mice (Fig. 2D-E). Expression of *Cxcl1* and *Cxcl2*, which act primarily to recruit neutrophils, were unchanged with misoprostol treatment. Given the established roles of CCL2 and CXCL5 in monocyte and neutrophil recruitment(31, 33), these data suggest that misoprostol dampens colonic inflammation by limiting chemokine-driven recruitment of these innate immune cells to the cLP.

**Fig. 2.**
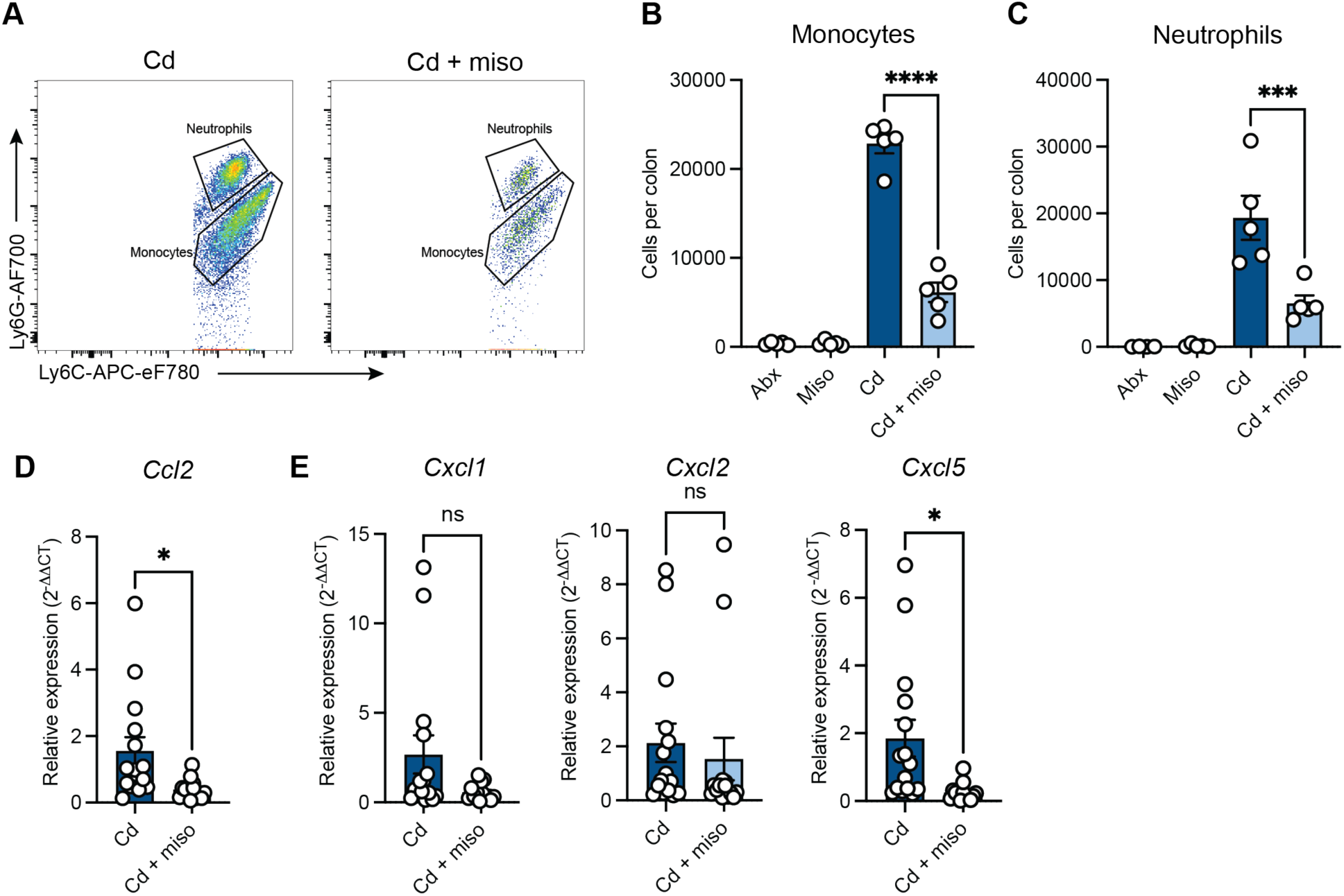
Misoprostol reduces colonic monocyte and neutrophil infiltration during CDI. (**A**-**E**) Mice were infected with *C. difficile* (Cd) and treated with or without misoprostol (miso). Samples were collected 2 days post-infection. (**A**) Representative flow cytometry plots of monocytes and neutrophils in the cLP. (**B** and **C**) Quantification of total (**B**) monocytes and (**C**) neutrophils in the cLP. Antibiotic (Abx) and misoprostol (Miso) treated uninfected controls (N = 5 per group). (**D** and **E**) RT-qPCR of (**D**) monocyte- and (**E**) neutrophil-recruiting chemokines in bulk colon tissue harvested from mice. Data relative to average *Gapdh* expression of Cd mice (N = 15 per group, representative of three independent experiments). (**B**-**E**) Each dot represents an individual mouse. Data are represented as mean ± SEM. Statistics by one-way ANOVA test with Tukey’s multiple comparisons (**B**-**C**) and unpaired t test (**D**-**E**). ns (not significant) P>0.05, *P<0.05, ***P<0.001, ****P<0.0001.

### Misoprostol alters systemic neutrophil levels during infection by reducing mobilization from the bone marrow

Our data demonstrate that misoprostol alters recruitment of critical innate immune cells to the gastrointestinal tract during infection. To begin to define the underlying mechanisms of misoprostol-mediated immunomodulation during infection, we asked if the effects of misoprostol treatment were restricted to the gastrointestinal tract. Using flow cytometry, we quantified monocyte and neutrophil frequencies in the blood two days post-infection(37) (Fig. S3B). We did not find any significant differences in the frequencies of circulating monocytes between the groups (Fig. 3A), supporting that misoprostol reduces colonic infiltration of monocytes through altered chemokine expression. However, we observed marked reductions in the frequencies of circulating neutrophils in the blood of Cd + miso mice (Fig. 3B). These data suggest that the effect of misoprostol on the innate immune response extends beyond the site of infection.

**Fig. 3.**
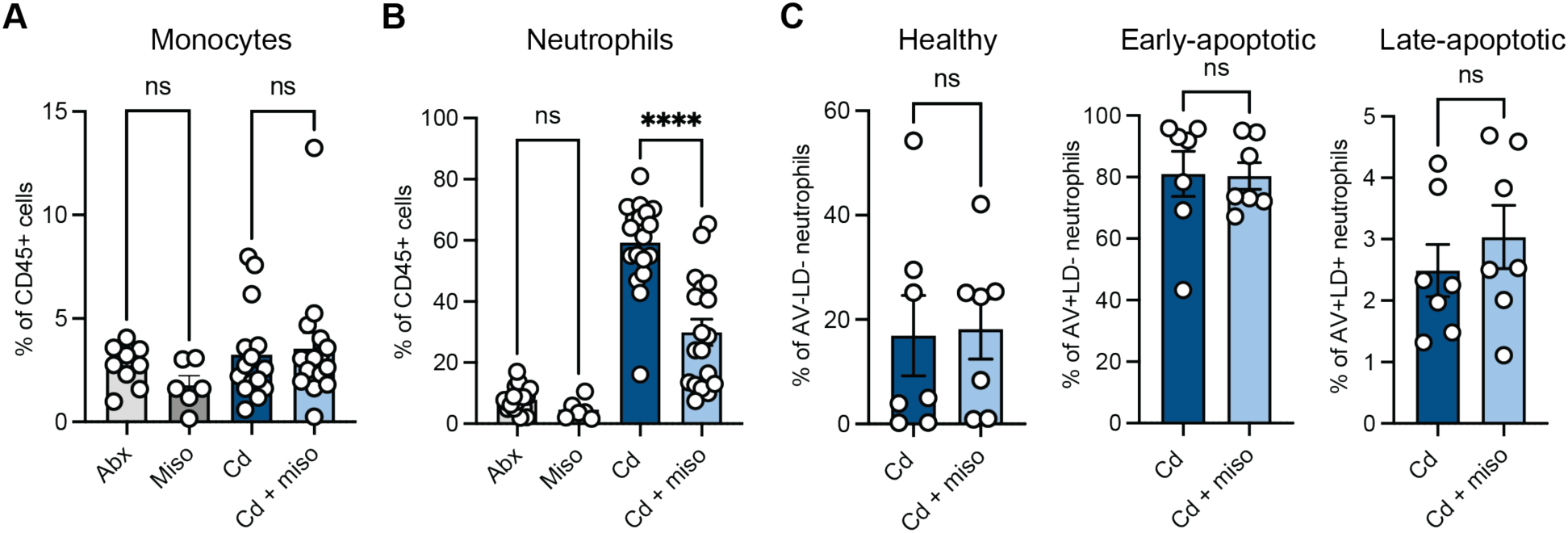
Misoprostol reduces systemic neutrophil levels during CDI. (**A**-**D**) Mice were infected with *C. difficile* (Cd) and treated with or without misoprostol (miso). Samples were collected 2 days post-infection. (**A** and **B**) Frequency of (**A**) monocytes and (**B**) neutrophils in the peripheral blood (N = 6 – 19 per group, representative of four independent experiments). (**C**) Annexin V (AV) and amine reactive dye (LD) staining in blood neutrophils. Percent of AV-LD- (healthy), AV+LD- (early-apoptotic), and AV+LD+ (late-apoptotic) neutrophils (N = 7 per group, representative of two independent experiments). (**A**-**C**) Each dot represents an individual mouse. Data are represented as mean ± SEM. Statistics by one-way ANOVA test with Tukey’s multiple comparisons (**A**-**B**) and unpaired t test (**C**). ns (not significant) P>0.05, ****P<0.0001.

To determine how misoprostol reduces circulating neutrophil levels during infection, we next explored whether misoprostol impacts neutrophil survival or development. Neutrophils are short lived cells primarily due to their ability to rapidly turn on apoptotic pathways that aim to balance tissue inflammation and homeostasis(38). To test the potential effects of misoprostol on neutrophil survival, we used Annexin V (AV) and amine reactive dye (LD) staining to discriminate healthy (AV-LD-), early-apoptotic (AV+LD-), and late-apoptotic (AV+LD+) neutrophils(39). We found no differences in viability or apoptotic states in blood neutrophils across treatment groups (Fig. 3C) suggesting that reduced circulating neutrophils are not due to altered survival.

Given these findings, we next investigated whether misoprostol impacts neutrophil development and expansion in the bone marrow. Neutrophils arise from self-renewing hematopoietic stem cells (HSCs), which give rise to all blood cell lineages(40). HSCs are defined as Lin^-^Sca1^+^c-Kit^+^ (LSK) cells and can differentiate into multipotent progenitors (MPPs), which are subdivided into distinct lineages. Among these, MPP3s are biased toward monocyte/granulocyte differentiation and give rise to granulocyte-monocyte progenitors (GMPs) and granulocyte progenitors (GPs), which can ultimately generate neutrophils(41). Therefore, to examine whether misoprostol impacts granulopoiesis during CDI, we analyzed these bone marrow progenitor populations at two days post-infection(42) (Fig. S4, Table S1). During CDI, total cellularity of the femoral bone marrow was unchanged between Cd and Cd + miso mice (Fig. 4A). However, Cd + miso mice exhibited a significant increase in total LSK cells and concurrent expansion of short-term HSCs and MPP3s (Fig. 4B), consistent with previous studies showing that PGE_2_ promotes HSC expansion(43-45). However, total numbers of downstream GMPs and GPs were unaffected (Fig. 4C), and there was no significant difference in total neutrophil numbers in the bone marrow between groups (Fig. 4D). These data suggest that while misoprostol influences early hematopoietic progenitors during CDI, the reduction in circulating neutrophils is likely not due to decreased production but instead due to altered egress from the bone marrow.

**Fig. 4.**
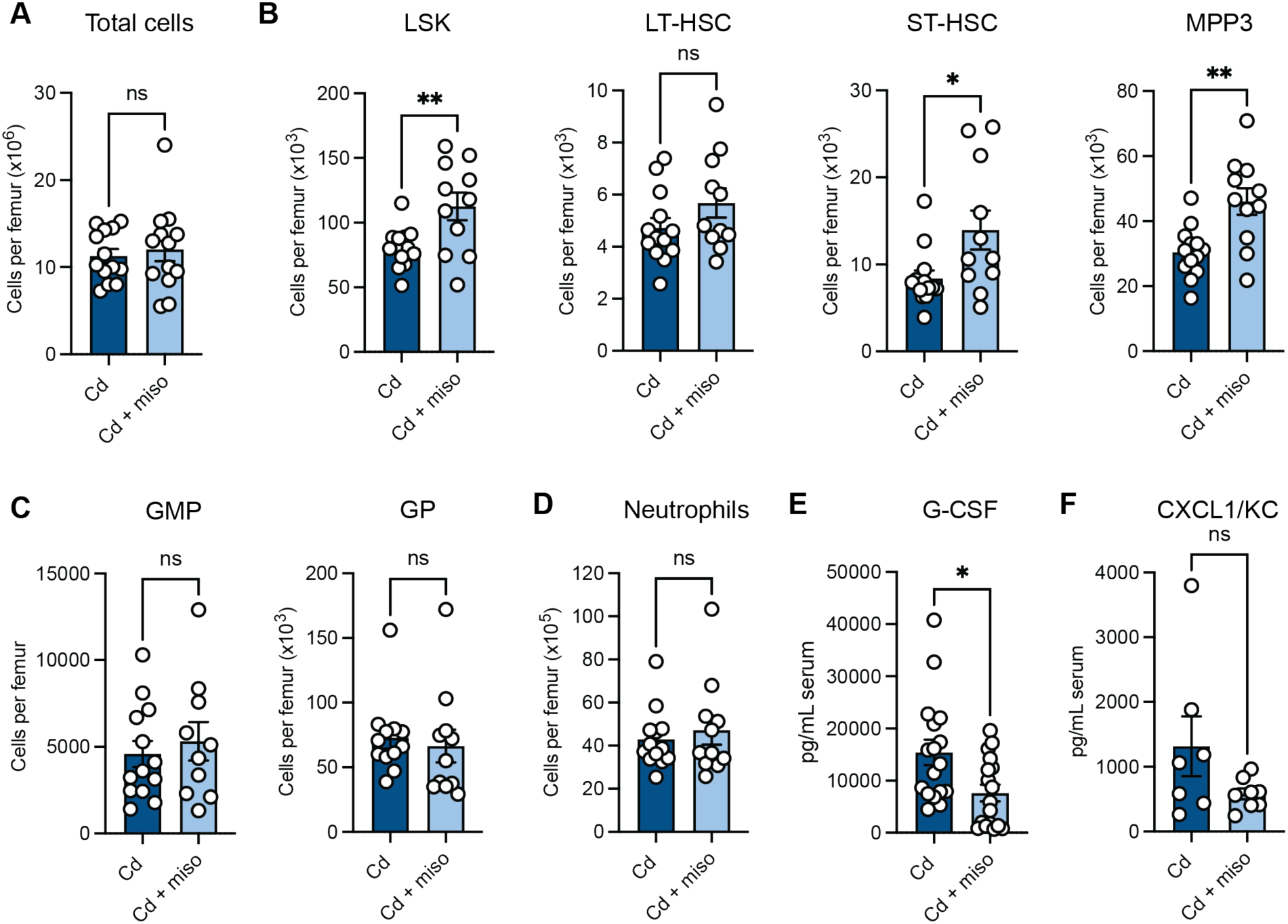
Misoprostol expands early hematopoietic progenitors without altering neutrophil development in the bone marrow. (**A**-**E**) Mice were infected with *C. difficile* (Cd) and treated with or without misoprostol (miso). Samples were collected 2 days post-infection. (**A**) Cellularity of femoral bone marrow (N = 13 per group). (**B**) LSK (Lin^-^Sca1^+^c-Kit^+^) cells per femur. These cells were further characterized as long-term HSCs (LT-HSC) (CD135^-^CD48^-^CD150^+^LSK), short-term HSCs (ST-HSC) (CD135^-^CD48^-^CD150^-^LSK), or multipotent progenitor 3 cells (MPP3) (CD135^-^ CD48^+^CD150^-^LSK) (N = 10 – 13 per group). (**C**) Granulocyte-monocyte progenitor (GMP) (Lin^-^ Ly6G^+^c-Kit^+^CD34^+^CD16/32^hi^Ly6C^-^CD135^-^CD115^-^) and granulocyte progenitor (GP) (Lin^-^Ly6G^+^c- Kit^+^CD34^+^CD16/32^hi^Ly6C^+^CD135^-^CD115^-^) cells per femur (N = 10 – 13 per group). (**D**) Neutrophils (Lin^+^Ly6G^+^Ly6C^+^) per femur (N = 11 – 13 per group). (**E**) Serum G-CSF levels quantified by ELISA (N = 17 per group). (**F**) Serum CXCL1/KC levels in mice measured by ELISA (N = 7 – 8 per group). (**A**–**F**) Each dot represents an individual mouse. Data are represented as mean ± SEM and representative of at least three independent experiments. Statistics by unpaired t test. ns (not significant) p>0.05, *P<0.05, **P<0.01.

Based on these observations, we hypothesized that misoprostol may impair mobilization of bone marrow neutrophils into circulation. Granulocyte colony-stimulating factor (G-CSF) is a critical regulator of granulopoiesis during both homeostasis and emergency granulopoiesis in response to infection(46). Along with its role in neutrophil development, G-CSF is also a potent inducer of neutrophil mobilization and release from the bone marrow into the bloodstream(26, 47, 48). Additionally, chemokines, such as CXCL1/KC, are critical for neutrophil recruitment from the bone marrow into circulation(33). To test whether misoprostol impacts this process, we measured circulating levels of serum G-CSF at two days post-infection by ELISA. We found that Cd + miso mice had significantly reduced serum G-CSF levels compared to Cd mice (Fig. 4E). Notably, we did not observe any differences in serum CXCL1/KC levels between groups (Fig. 4F). Previous studies have shown that mice with impaired G-CSF signaling exhibit decreased circulating neutrophils despite normal levels of mature neutrophils in the bone marrow(26). Collectively, these data suggest that during CDI, misoprostol may limit neutrophil mobilization from the bone marrow by dampening G-CSF production, resulting in decreased circulating neutrophils.

### Pharmacological modulation of neutrophil infiltration tunes disease severity during CDI

Our data suggest that misoprostol reduces CDI disease severity by reducing circulating neutrophils and recruitment to the site of infection through modulation of G-CSF levels and chemokine expression. Because G-CSF promotes neutrophil mobilization into circulation(26, 47, 48), we hypothesized that G-CSF supplementation would restore blood neutrophils and counteract the effects of misoprostol. To test this, we treated mice with misoprostol and 250μg/kg recombinant mouse G-CSF (rG-CSF) daily beginning one day prior to infection with *C. difficile* (Cd + miso + rG-CSF) (Fig. 5A). We observed that rG-CSF treatment restored neutrophil frequencies in the blood and cLP in Cd + miso + rG-CSF mice to those observed in Cd mice at two days post-infection (Fig. 5B). Additionally, rG-CSF treatment did not impact monocyte frequencies (Fig. S2A) or expression of monocyte and neutrophil chemokines in the colon (Fig. S2B-C). Notably, this treatment and concurrent increases in systemic and colonic neutrophil levels significantly increased severity of disease following infection. However, disease was not as severe as Cd mice (Fig. 5C), suggesting that misoprostol reduces CDI severity in part by limiting neutrophil infiltration but also exerts other protective effects that further mitigate disease.

**Fig. 5.**
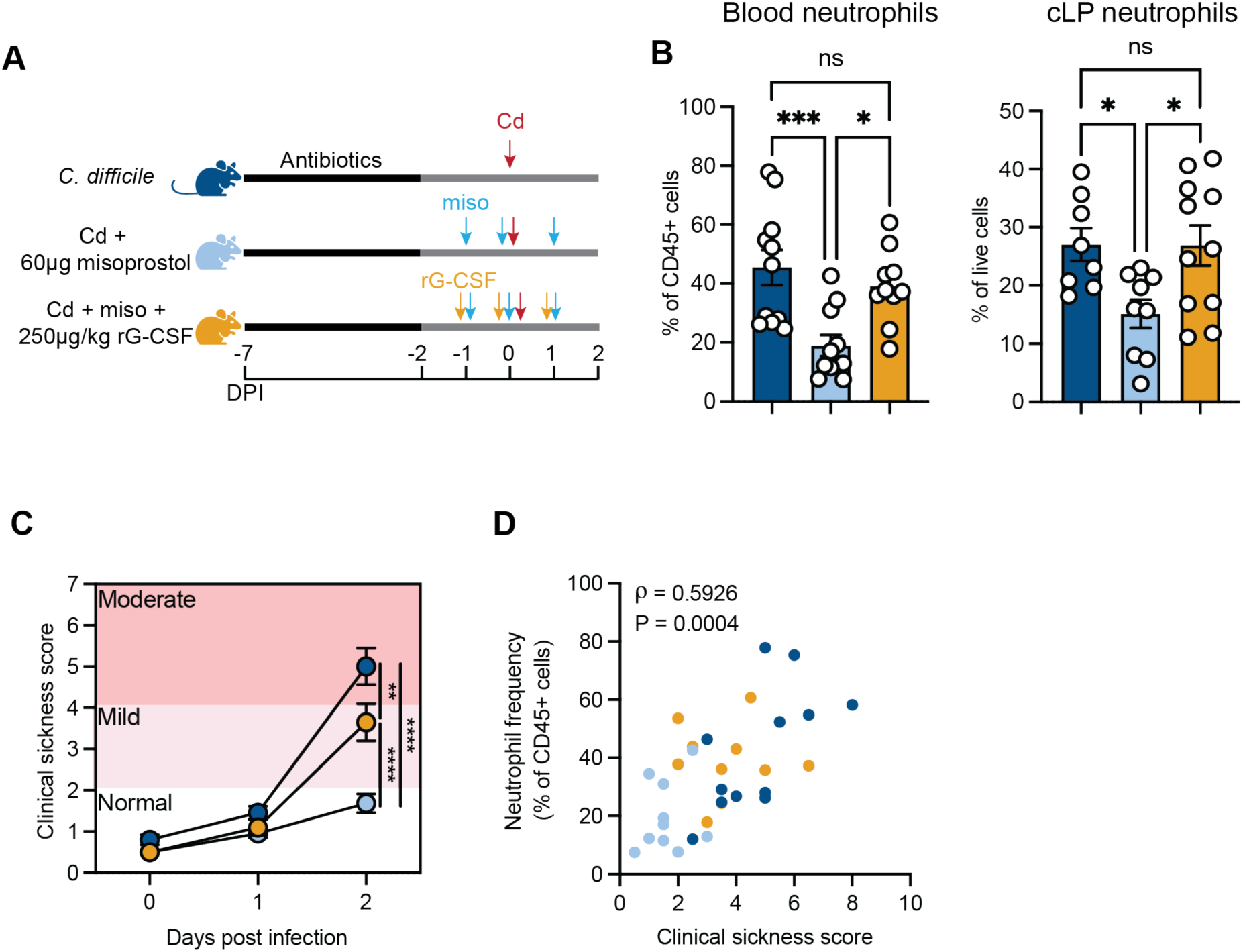
Pharmacological neutrophil modulation tunes CDI severity. (**A**) Schematic of experimental design. Mice were treated with antibiotics (cefoperazone) in their drinking water and infected with 1x10^5^ *C. difficile* spores (Cd). Misoprostol-treated mice were administered 60 μg misoprostol via intraperitoneal (i.p.) injection daily from -1 days post-infection (DPI) to 1 DPI. rG- CSF-treated mice received 250 μg/kg mouse recombinant G-CSF via i.p. injection daily concurrently with misoprostol. (**B**) Frequency of neutrophils in the peripheral blood and cLP at 2 DPI (N = 8 – 11 per group). (**C**) Composite score of weight loss, stool consistency, and behavior (N = 10 – 11 per group). (**D**) Two-sided Spearman correlation between blood neutrophil frequency and clinical sickness score of *C. difficile-*infected mice at 2 DPI. Each dot represents an individual mouse and is color-coded by treatment group (Spearman *π* = 0.5926, N = 32). (**B**–**C**) Data are represented as mean ± SEM and representative of two independent experiments. Statistics by one-way (**B**) or two-way (**C**) ANOVA with Tukey’s multiple comparisons. ns (not significant) P>0.05, *P<0.05, **P<0.01, ***P<0.001, ****P<0.0001.

Finally, to contextualize these data and assess the relationship between neutrophils and *C. difficile*-associated disease, we examined correlations between circulating neutrophil frequencies in the blood and CDI severity. Across infected mice, circulating neutrophils positively correlated with disease severity. Strikingly, reduced neutrophil frequencies were associated with decreased severity in Cd + miso mice. Additionally, misoprostol and rG-CSF treatment restored both neutrophil levels and severity to intermediate levels between Cd + miso and Cd mice (Fig. 5D). Together, these findings underscore the pivotal role of neutrophil abundance in driving CDI pathology and highlight the potential of modulating innate immune responses as a treatment strategy to combat CDI.

## DISCUSSION

In this study, we identify a novel mechanism by which the innate immune system can be modulated to improve CDI outcomes. CDI remains difficult to treat with high recurrence rates and increasing antibiotic resistance. Although antibiotics are used to treat infections, they perpetuate microbiota disruption and do not address the host inflammatory responses that strongly correlate with disease severity(4, 5). We previously reported that misoprostol improves CDI survival in mice, but the mechanism of protection remained unclear(17). Here, we demonstrate that misoprostol reduces CDI severity by limiting G-CSF production, thereby reducing neutrophil availability and subsequent infiltration into the colon.

PGs have key roles in intestinal homeostasis, intestinal wound repair, and mucosal protection, but their function in CDI has not been extensively explored(15). We found that misoprostol reduces epithelial injury during CDI (Fig. 1F). However, we observed no significant differences in IEC proliferation, inflammatory responses, or mucus-related gene expression at two days post-infection (Fig. S1), in contrast to previous studies, which have shown that PGE_2_ can enhance IEC proliferation, mucus secretion, and barrier protection in the context of small intestinal enteropathy and inflammatory bowel disease.(23-25) Notably, misoprostol did increase IEC proliferation in uninfected mice (Fig. S1E), suggesting that earlier or transient effects on the epithelium prior to CDI could contribute to barrier fortification upon infection. However, further work will be needed to further test this hypothesis.

Our data highlight immunomodulation as the dominant mechanism of misoprostol-mediated protection. E-series PGs, like PGE_2_, have been shown to dampen innate immune cell activation and proinflammatory cytokine production during infection(16, 19, 36, 49). Consistent with this, misoprostol reduced colonic *Tnfa* expression and levels of the neutrophil-derived inflammatory marker calprotectin (Fig. 1G-H). Notably, misoprostol also reduced total neutrophils in the lamina propria (Fig. 2C) and neutrophil frequency in the blood (Fig. 3B), which were previously unreported. Misoprostol also decreased expression of chemokines that drives neutrophil recruitment (Fig. 2E). Since chemokine gradients orchestrate neutrophil trafficking, suppressing these signals may help limit intestinal infiltration. Neutrophils are a hallmark of CDI and correlate with greater disease severity(5), yet the mechanisms linking neutrophil influx to tissue damage during CDI remain incompletely defined. Neutrophils produce high levels of inflammatory molecules, including TNFα and reactive oxygen species(19). In other enteric infection models, PGE_2_ has been shown to reduce neutrophil activation and function, resulting in decreased TNFα and ROS production(19). These findings suggest that misoprostol may similarly attenuate neutrophil function during CDI, limiting additional inflammation and damage. Additionally, excessive recruitment of neutrophils to the colonic mucosa can cause crypt abscesses and transepithelial migration that lead to increased intestinal barrier permeability(50, 51). Hence, a reduction in neutrophil infiltration and potentially activation, likely contributes to the decrease in colonic inflammation and reduced epithelial tissue damage we observe.

PGE_2_ has a well-established role in hematopoiesis, promoting stem cell proliferation and survival(44, 45). Here, we found that misoprostol, a PGE_2_ mimetic, phenocopies these effects, increasing the total number of HSCs and reshaping the early immune progenitor pool, expanding ST-HSCs and MPP3s as seen with PGE_2_. Surprisingly, misoprostol treatment did not alter total numbers of later neutrophil progenitors or bone marrow neutrophils (Fig. 4A-D). This suggests that misoprostol’s impact on granulopoiesis is unlikely to directly account for the reduced neutrophils observed in the blood and colon. Nevertheless, its effects on early progenitors could influence neutrophil production at different stages of infection by potentially supporting survival and preventing stem cell exhaustion or altering the differentiation potential of these cells and thereby enabling more balanced or sustained neutrophil outputs. However, future studies will be needed to test these hypotheses.

G-CSF is a key cytokine that drives neutrophil development and mobilization during emergency granulopoiesis(26, 48). Here, we show that Cd + miso mice had reduced circulating G-CSF (Fig. 4E), potentially limiting neutrophil mobilization from the bone marrow. To test whether restoring G-CSF could overcome this effect during CDI, we administered rG-CSF to misoprostol-treated mice, resulting in increased circulating neutrophils and, unexpectedly, also enhanced neutrophil infiltration into the colon (Fig. 5B). Despite these increases, disease severity of Cd + miso + rG-CSF mice only reached an intermediate level between Cd + miso and Cd mice (Fig. 5C), highlighting that misoprostol’s protective effects extend beyond reducing neutrophil numbers and reflect broader anti-inflammatory actions. Strikingly, we observed a positive correlation between higher circulating neutrophil frequencies and increased CDI severity (Fig. 5D), supporting the concept that excessive neutrophil recruitment exacerbates pathology. Additional studies will be necessary to further understand how misoprostol impacts G-CSF production and subsequent effects on neutrophil mobilization. PGs have been shown to enhance CXCL12/CXCR4 signaling to retain hematopoietic cells in the bone marrow(44, 52), whereas G-CSF reduces CXCR4 expression to promote mobilization(47). Whether this axis contributes to our misoprostol phenotype remains unknown and represents an intriguing area for further study.

Collectively, this work identifies a novel mechanism by which misoprostol protects against CDI and provides a new framework for understanding how neutrophils contribute to CDI severity. By demonstrating that modulation of neutrophil mobilization and infiltration can markedly alter CDI outcomes, this study emphasizes the balance of neutrophil biology during CDI and establishes this innate immune cell type as a potential therapeutic target for host-directed therapy. Following our previous study, misoprostol was advanced into a clinical study to test its ability to reduce recurrent CDI(53). Although this study was limited by enrollment and retention challenges during the COVID-19 pandemic, our mechanistic data provide renewed support for evaluating misoprostol, or other neutrophil-modifying treatments, in clinical settings. Together, these findings highlight neutrophil infiltration as a critical driver of CDI pathology and establish misoprostol as a promising host-directed therapeutic strategy. While additional studies are needed to define its effects in humans, this work provides a foundation for developing targeted interventions that complement current treatments for CDI.

## MATERIALS AND METHODS

### Animal and experimental models of *C. difficile* infection

All mouse procedures were approved by the Children’s Hospital of Philadelphia Institutional Animal Care and Use Committee (IACUC). 5-week-old C57BL/6 male mice were purchased from Jackson Laboratories and infected with *C. difficile* as previously described. Mice were given one week to acclimate to their new environment. 0.5 g/L cefoperazone was administered in drinking water *ad libitum* for 5 days, followed by a 2-day recovery period before subsequent infection. Mice were administered 1x10^5^ spores of *C. difficile* CD196 in PBS via oral gavage. Mice were monitored daily for survival, weight loss, and clinical sickness(27). Mice were euthanized when they appeared moribund or weight loss exceeded 20% of their original starting weight.

### Misoprostol and rG-CSF treatment

Misoprostol was purchased from Cayman Chemicals (Ann Arbor, MI) in methyl acetate. The solvent was evaporated under nitrogen, and the misoprostol was resuspended in PBS immediately before each administration. 60 μg misoprostol was administered daily via intraperitoneal (i.p.) injection starting one day prior to infection.

Mouse G-CSF recombinant protein (rG-CSF) was purchased from PeproTech and resuspended in water according to manufacturer’s instructions. 250 μg /kg rG-CSF was administered via i.p. injection daily starting one day prior to infection.

### *C. difficile* burdens and toxin titers from feces

For quantification of *C. difficile* burdens, fecal samples were weighed, homogenized in sterile PBS, and plated on taurocholate cycloserine cefoxitin fructose agar (TCCFA) under anaerobic conditions. *C. difficile* toxin titers were quantified using a Vero cell rounding assay as previously described(54). Briefly, homogenized fecal samples were centrifuged, and the supernatant was filtered through a 0.2 μm filter. Supernatants were diluted and incubated overnight with Vero cell monolayers. Toxin titers were calculated as the reciprocal value of the highest dilution with 100% cell rounding.

### Histological analysis

Samples were fixed in 10% formalin and embedded in paraffin as previously described(7). Slides were stained with Hematoxylin and Eosin and assigned a disease score by a blinded pathologist based on previously described criteria(55). Histological scores are reported as a cumulative score of three independent criteria: epithelial damage, inflammation, and edema.

Ki67 staining was performed as previously described(56). The immunohistochemistry slides were scanned x40. Using QuPath5.1, an object classifier was built to distinguish epithelial cells from immune cells and other cells using a random forest classifier after numerous annotations from an image created from samples of 15 case images. The percentage of KI67 positive epithelial cells was determined by deploying the positive cell detection and built object classifier tools(57).

### Fecal and serum ELISAs

Lipocalin-2 and calprotectin were quantified from stool using mouse Lipocalin-2/NGAL and S100A8/S100A9 R&D DuoSet ELISA kits according to manufacturer’s instructions. Fecal pellets were homogenized in 1ml sterile PBS and centrifuged at 4000rpm for 5 minutes. Supernatants from *C. difficile* or DSS-treated mice were diluted 1:100 for Lipocalin-2 and 1:50 for calprotectin quantification. Protein concentrations were normalized to grams of feces.

CXCL1/KC and G-CSF were quantified from serum using R&D DuoSet ELISA kits according to manufacturer’s instructions. Blood was collected via cardiac puncture, allowed to coagulate, and centrifuged at 8000g for 10min. Serum was collected and diluted 1:10 for CXCL1/KC and 1:30 or 1:50 for G-CSF quantification.

### RNA extraction and RT-qPCR

RNA was extracted using RNeasy Mini Kit (QIAgen) according to manufacturer’s instructions. cDNA was synthesized using M-MLV Reverse Transcriptase (Promega) according to manufacturer’s instructions. RT-qPCR was performed using iQ SYBR reagents (Bio-rad), and data were normalized to expression of *Gapdh*. The primer sequences were used as follows: *Ccl2*, GCTACAAGAGGATCACCAGCAG (forward) and GTCTGGACCCATTCCTTCTTGG (reverse); *Cxcl1*, TCCAGAGCTTGAAGGTGTTGCC (forward) and AACCAAGGGAGCTTCAGGGTCA (reverse); *Cxcl2*, CATCCAGAGCTTGAGTGTGACG (forward) and GGCTTCAGGGTCAAGGCAAACT (reverse); *Cxcl5*, CCGCTGGCATTTCTGTTGCTGT (forward) and CAGGGATCACCTCCAAATTAGCG (reverse); *Tnfa*, GCCTCTTCTCATTCCTGCTTG (forward) and CTGATGAGAGGGAGGCCATT (reverse); *Muc2*, AAACTGCTCTCTGGACTGCC (forward) and TTGGTTGGTGTGCTGAGTGT (reverse); *Gapdh*, AGCAAGGACACTGAGCAAGAG (forward) and GCAGCGAACTTTATTGATGGT (reverse).

### Isolation of cells for flow cytometry

Intestinal epithelial cells (IECs) and colonic lamina propria cells (cLPs) were isolated from mice as previously described(58). Briefly, colons and ceca were collected, opened longitudinally, washed in PBS, and cut into 1cm pieces. The tissue was incubated in HBSS with 2mM EDTA and shaken for 10 minutes at 150 rpm to dissociate IECs. Supernatants containing IECs were filtered through 70μm cell strainer and used for RT-qPCR. Remaining tissue pieces were washed, minced, and incubated in digestion buffer containing RPMI 1640, 5% FBS, 0.5 mg/mL collagenase type VIII, 5 U/mL DNase, 100 IU/mL penicillin and 100 μL/mL streptomycin for 45 minutes at 37°C. Cell suspensions were filtered through 70μm cell strainer and stained directly.

Blood was obtained via cardiac puncture and collected in heparin containing tubes. Following red blood cell lysis with ACK lysis buffer, cells were washed and pelleted at 400g for 5 minutes. ACK lysis was repeated until cell pellets were no longer red. Cells were enumerated and transferred to U-bottom 96-well plate for staining.

Both femurs were collected from mice for bone marrow isolation as previously described(59). Briefly, connective tissue was removed by gentle scraping with a razor before one end of each femur was cut to expose the marrow. The femurs were then transferred into a 0.5ml microcentrifuge tube with a hole punctured in the bottom with an 18g needle. The 0.5ml tube was then placed in a 1.5ml tube and centrifuged at 13,000g for 1.5min. Cell pellets were then lysed in ACK buffer, washed, and pelleted at 400g for 5min before being passed through at 40 μm filter and resuspended in PBS. Cells were enumerated and equivalent numbers of cells were transferred into a U-bottom 96-well plate for staining.

### Flow cytometry and antibodies

Cell suspensions were prepared as described above and stained with LIVE/DEAD Fixable Aqua (1:600) or Fixable Viability Dye 660 (1:1000) in PBS. Cells were then suspended in Fc block (5ug/ml CD16/32 in FACS buffer). If CD16/32 was stained for, the staining antibody was substituted for the unconjugated antibody.

For phenotypic surface analysis of cLP monocytes and neutrophils, cells were stained as previously described(19) with antibodies against CD3-PerCP-eF710 (17A2, Invitrogen) and B220-PerCP/Cy5.5 (RA3-6B2, eBioscience) at 1:200 dilution, CD11b-PE-Texas Red (M1/70.15, Invitrogen) and Ly6G-AF700 (1A8, Biolegend) at 1:300, and Ly6C-APC-eF780 (HK1.4, eBioscience) at 1:800 in FACS buffer.

To quantify blood monocytes and neutrophils, cells were stained with antibodies against CD45-APC/Cy7 (clone 30-F11, BD Biosciences), CD11b-V450 (clone M1/70, BD Biosciences), Ly6G-PerCP/Cy5.5 (Clone 1A8, CD Biosciences) and Ly6C-FITC (HK1.4, Biolegend) at 1:150 dilution in FACS buffer. To further assess neutrophil viability, cells were stained with antibodies against Annexin V-PE/Cy7 (1:20, Invitrogen) according to manufacturer’s instructions. Early-apoptotic neutrophils were defined as Annexin V+ Fixable Viability Dye 660-, and late-apoptotic neutrophils were defined as Annexin V+ Fixable Viability Dye 660+.

For bone marrow progenitor surface profiling, cells were stained with antibodies against CD3ε-APC (145-2C11, Biolegend), CD11b-APC (M1/70, Biolegend), CD45R/B220-APC (RA3-6B2, Biolegend), TER-119-APC (Biolegend), CD115-BV711(AFS98, Biolegend), Sca1-Pacific Blue (E13-161.7, Biolegend), c-Kit-BV785 (2B8, Biolegend), and CD48-APC/Cy7 (HM48-1, Biolegend) at 1:200 dilution, CD150-PerCP/Cy5.5 (TC15-12F12.2, Biolegend), CD34-FITC (RAM34, eBioscience), CD135-BV421 (A2F10, Biolegend) at 1:100, and Ly6C-AF700(HK1.4, Biolegend) and Ly6G-BUV563 (1A8, BD) at 1:300 in FACS buffer. Samples were run on a Cytek Aurora and data were analyzed using FlowJo software.

### Statistical analysis

Statistical analyses were performed using GraphPad Prism version 10.6.0. Specific tests are reported in the figure legends.

## ACKNOWLEDGEMENTS

The authors would like to thank all members of the Zackular laboratory for providing feedback and support during this study. We thank the CHOP Flow Core for support.

## Funding

This work was supported by:

Children’s Hospital of Philadelphia’s Center for Microbial Medicine

National Institutes of Health grant U19AI174998-03 (JPZ)

National Institutes of Health grant R01AI187174-01A1 (JPZ)

National Institutes of Health grant R01AI148249 (CAH)

## Author contributions

Conceptualization: OK, JSO, AS, JPZ

Methodology: OK, JSO, AS, EEF, GCWB, DLA

Investigation: OK, JSO, AS, THZ, EEF

Visualization: OK, JPZ

Supervision: OK, JPZ

Writing – original draft: OK, JPZ

Writing – reviewing and editing: OK, JSO, AS, THZ, GCWB, DLA, CAH, KC, DMA, JPZ

## Data and materials availability

All data needed to evaluate the conclusions in the paper are available in the main text or the supplementary materials.

---

**Fig. S1.**
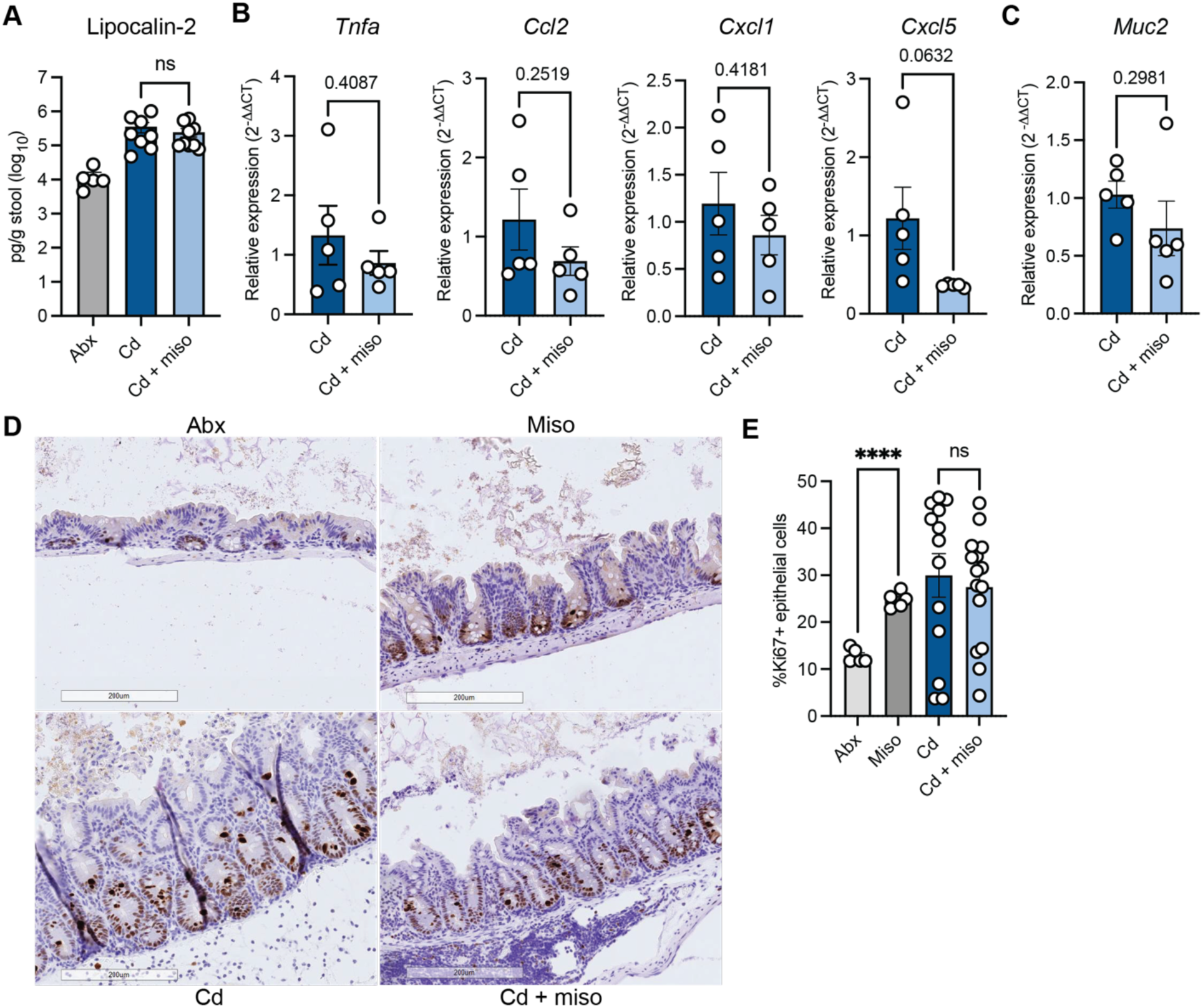
Misoprostol-mediated protection is not associated with the intestinal epithelial response to infection. (**A**) Fecal lipocalin-2 levels of antibiotic-treated uninfected controls (Abx,), *C. difficile-*infected mice (Cd), and *C. difficile*-infected and misoprostol-treated mice (Cd + miso) at two days post-infection (DPI) measured by ELISA (N = 5 – 9 per group). (**B** and **C**) RT-qPCR of (**B**) proinflammatory cytokine and chemokine and (**C**) mucin-related gene expression in IECs harvested from mice at 2 DPI. Data relative to average *Gapdh* expression of Cd mice (N = 5 per group). (**D**) Representative images of Ki67 staining in ceca from antibiotic-treated (Abx), misoprostol-treated (Miso), *C. difficile* infected (Cd), and *C. difficile* infected and misoprostol-treated mice (Cd + miso). (**E**) Quantification of Ki67+ epithelial cells (N = 5 – 15 per group, representative of three independent experiments). (**A**–**E**) Data are represented as mean ± SEM. Statistics by one-way ANOVA test with Dunnett’s T3 multiple comparisons (**A**, **E**) and unpaired t test (**B**-**C**). ns (not significant) P>0.05, ****p<0.0001.

**Fig. S2.**
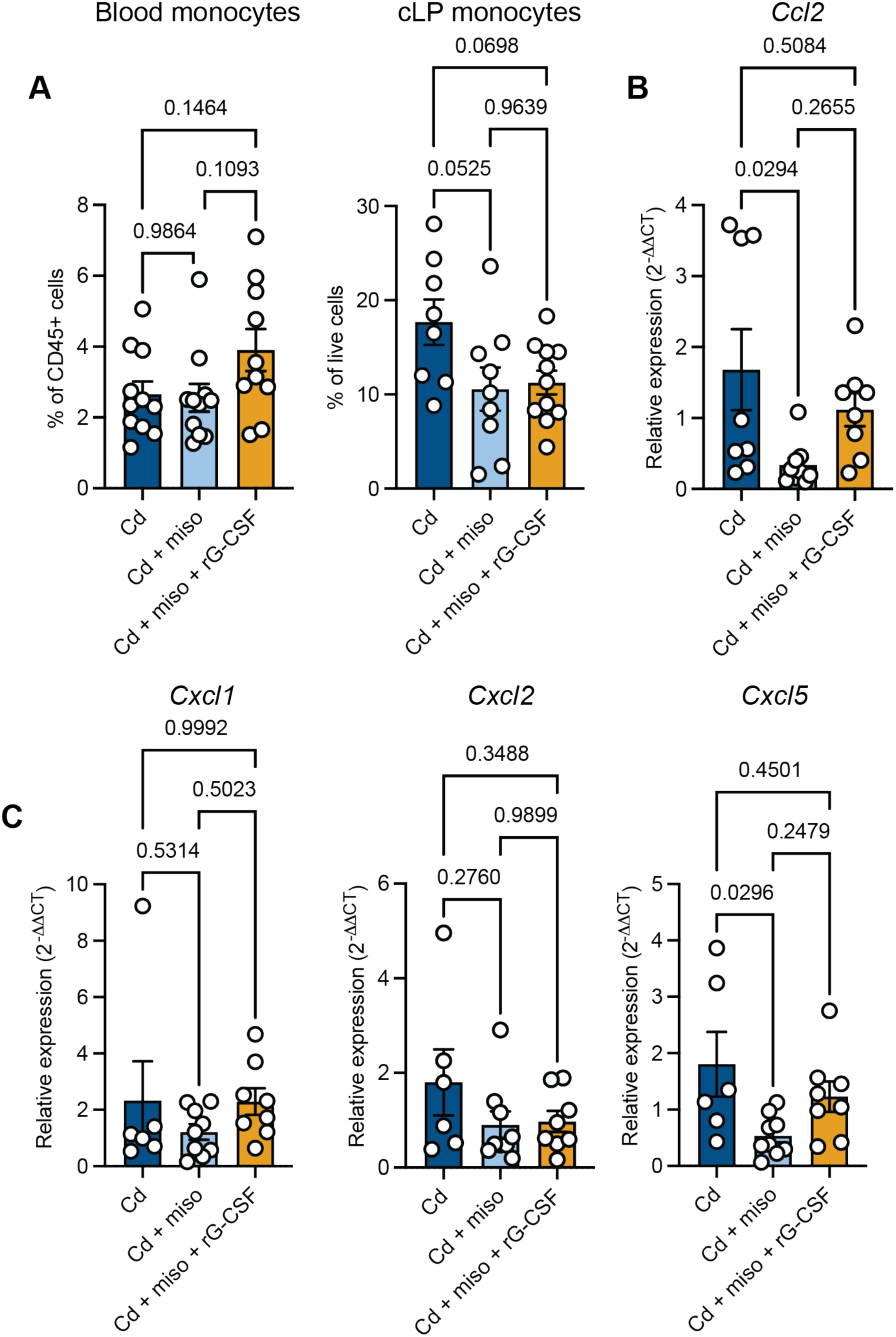
rG-CSF treatment does not impact monocyte frequencies or colonic chemokine expression during CDI. (**A**) Frequency of monocytes in the peripheral blood and cLP at 2 DPI (N = 8 – 11 per group). (**B** and **C**) RT-qPCR of (**B**) monocyte- and (**C**) neutrophil-recruiting chemokines in bulk colon tissue harvested from mice. Data relative to average *Gapdh* expression of Cd mice (N = 6 – 9 per group, representative of three independent experiments). (**A**-**C**) Data are represented as mean ± SEM and representative of two independent experiments. Statistics by one-way ANOVA with Tukey’s multiple comparisons.

**Fig. S3.**
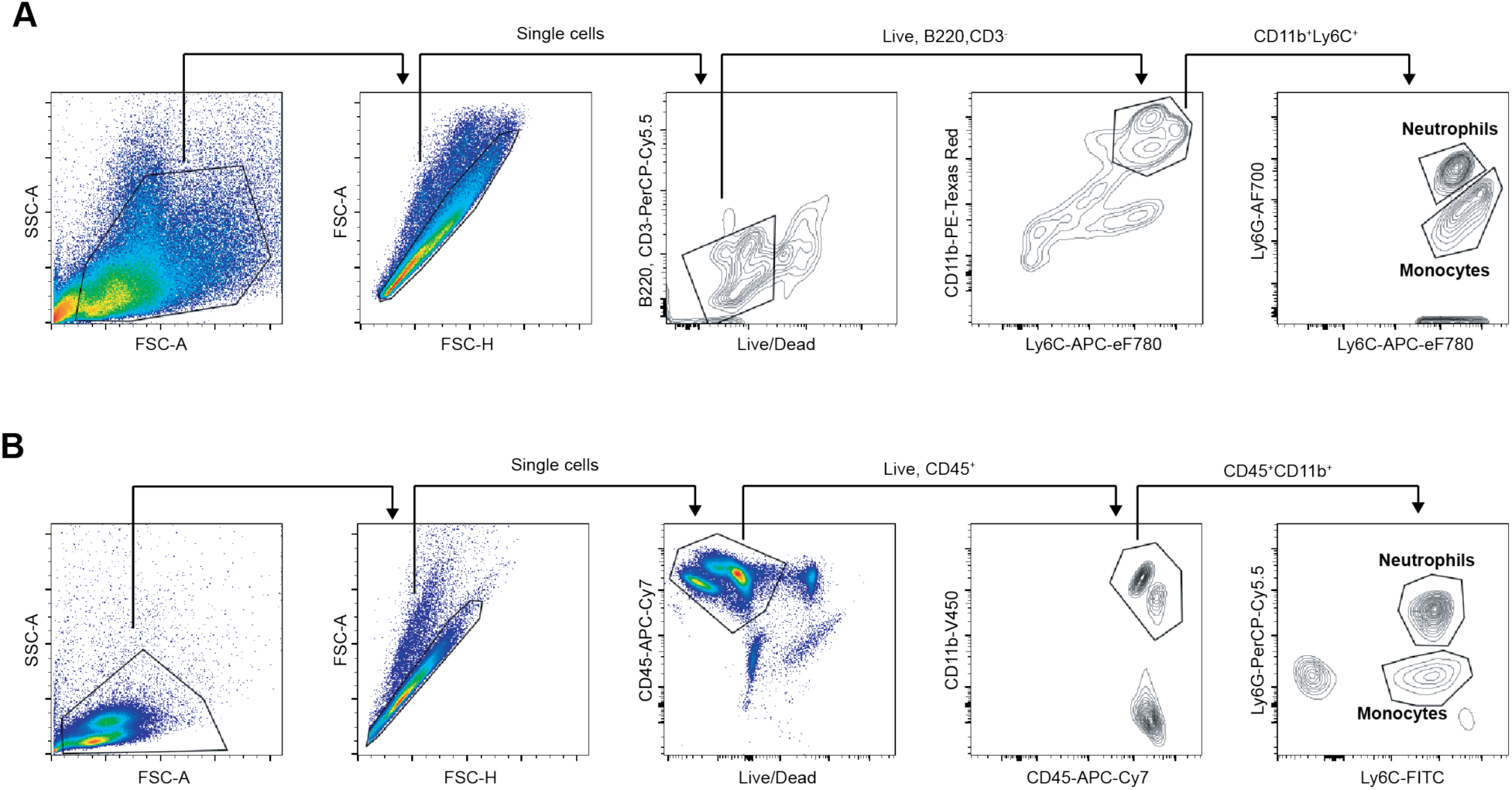
Gating of neutrophils and monocytes in the cLP and blood. Representative flow plots from the (**A**) cLP or (**B**) blood are shown indicating the gating strategy used to identify neutrophils and monocytes.

**Fig. S4.**
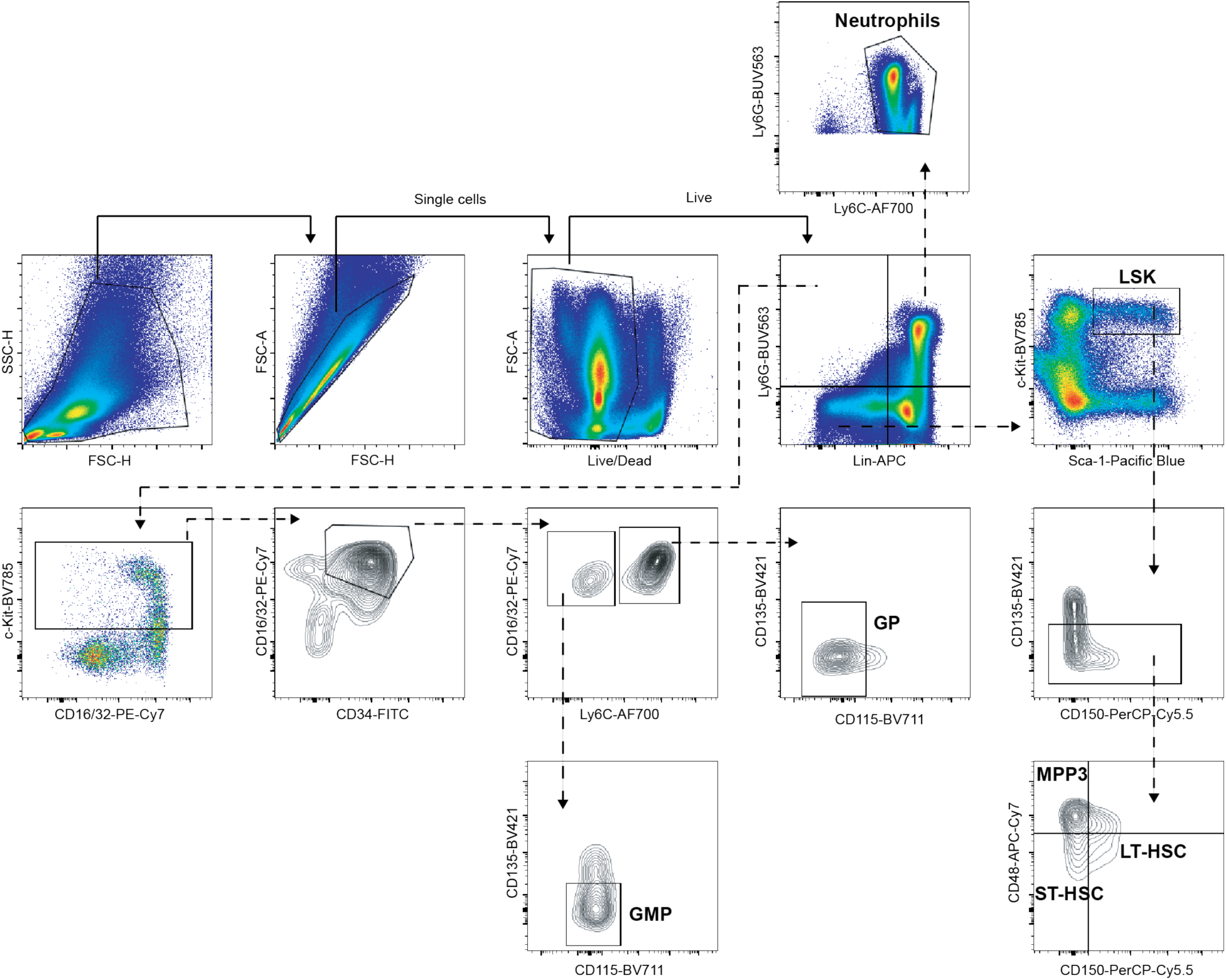
Gating of HSCs and neutrophil progenitors in the bone marrow. Representative flow plots from the bone marrow are shown indicating the gating strategy used to identify various cell populations.

**Table S1.** Surface markers of cells identified in the bone marrow.

| Cell type | Surface Markers |
| --- | --- |
| LSK | Lin(CD3ε, CD11b, CD45R/B220, TER119)<sup>-</sup>,<br>Ly6G<sup>-</sup>, Sca-1<sup>+</sup>, c-Kit<sup>+</sup> |
| Long-term hematopoietic stem cell (LT-HSC) | LSK, CD135<sup>-</sup>, CD48<sup>-</sup>, CD150<sup>+</sup> |
| Short-term hematopoietic stem cell (ST-HSC) | LSK, CD135<sup>-</sup>, CD48<sup>-</sup>, CD150<sup>-</sup> |
| Multipotent progenitor 3 (MPP3) | LSK, CD135<sup>-</sup>, CD48<sup>+</sup>, CD150<sup>-</sup> |
| Granulocyte-monocyte progenitor (GMP) | Lin(CD3ε, CD11b, CD45R/B220, TER119)<sup>-</sup>,<br>Ly6G<sup>+</sup>, c-Kit<sup>+</sup>, CD16/32<sup>hi</sup>, CD34<sup>+</sup>, Ly6C<sup>-</sup>,<br>CD135<sup>-</sup>, CD115<sup>-</sup> |
| Granulocyte progenitor (GP) | Lin(CD3ε, CD11b, CD45R/B220, TER119)<sup>-</sup>,<br>Ly6G<sup>+</sup>, c-Kit<sup>+</sup>, CD16/32<sup>hi</sup>, CD34<sup>+</sup>, Ly6C<sup>+</sup>,<br>CD135<sup>-</sup>, CD115<sup>-</sup> |
| Neutrophil | Lin(CD3ε, CD11b, CD45R/B220, TER119)<sup>+</sup>,<br>Ly6G<sup>+</sup>, Ly6C<sup>+</sup> |

